# Pretreatment with estrogen enhances the therapeutic efficacy of cardiac progenitor cells and improves cardiac recovery in a failing heart model

**DOI:** 10.1101/2025.06.25.661574

**Authors:** Dunya Aydos, Neslihan Basak, Yusuf Olgar, Kardelen Genc, Recep Uyar, Okan Ekim, Taras Sych, Zeynep Busra Aksoy, Tugba Aktan Kosker, Erdinc Sezgin, Ceylan Verda Bitirim

**Affiliations:** Ankara University, Stem Cell Institute, Ankara, Türkiye; Ankara University Faculty of Medicine Department of Biophysics, Ankara, Türkiye; Johannes Gutenberg University, Mainz, Germany; Veterinary Faculty, Ankara University, Ankara, Türkiye; Science for Life Laboratory, Department of Women’s and Children’s Health, Karolinska Institutet, Tomtabodavagen 23, 17165, Solna, Sweden

**Author notes:** **Correspondance:** Ceylan Verda Bitirim.

## Abstract

**Background:** In recent years, cardiac progenitor cells (CPCs) are considered as a potential source of cell therapy for the treatment of heart failure. Despite the encouraging results of clinical trials showing that transplantation of CPCs improves function of infarcted hearts, reduced cell survival and inefficient engraftment into host tissue are still challenging issues.

**Methods:** CPCs were isolated from a male mouse heart and incubated with estrogen (10^-7^M for 48 hours) in vitro. The heart failure model was generated by intraperitoneally isoproterenol (ISO) treatment (200mg/kg for 6 days) in the female mouse. Either estrogen pretreated-CPC (E2-CPC) or untreated-CPCs (Control-CPC) were intramyocardially transplanted into the failure heart. Engraftment of transplanted cells were visualized by in vivo imaging, Y-chromosome staining (by Fluorescence In Situ Hybridization) and SRY gene for expression analysis. Cardiac functions were determined by analyzing electrophysiological changes. Pathological changes following the transplantation were also examined at the molecular level by immunostaining and western blotting from isolated heart samples and on heart tissue sections to address the efficiency of the E2-CPC transplantation. In addition to regenerative outcomes at animal level, the enhancing effect of estrogen on the therapeutic potential of CPC was examined through changes in migration, proliferation and mitochondrial energetics by in vitro experiments. The effects of estrogen in transcriptome profiling was also evaluated by RNA sequencing (RNA-seq) in CPCs.

**Results:** In vivo results demonstrated that estrogen pretreatment increased the retention rate of transplanted CPCs in failing heart. E2-CPC transplantation enhanced cardiac function and ameliorated pathological cardiac remodelling via inducing revascularization, proliferation while attenuating collagen formation and hypertrophy in failing heart. Our in vitro results have shown that estrogen treatment promotes migration, angiogenesis and proliferation capacities and adenosine triphosphate (ATP) production of CPCs *in vitro.* RNA sequencing provided further evidence that the change in the transcriptome profile of CPCs upon estrogen treatment improved their ability to cause reverse cardiac remodelling in the failing heart after transplantation.

**Conclusion:** Our data indicated that estrogen-pretreatment significantly improves cardiac recovery in a failing heart through enhancing efficacy of CPC transplantation. This study also emphasized underyling mechanisms of the improvement in function in CPC-based therapies which remain poorly understood.

## Background

Cardiovascular diseases (CVDs) are considered one of the leading causes of morbidity and mortality worldwide, even with developments in modern surgical and medical techniques. Cardiovascular diseases, such as ischemic heart disease and cardiomyopathy, are associated with a lack of function in cardiomyocytes and/or a loss of healthy cardiomyocyte numbers [1]. Although the damage in the heart tissue induced by small infarcts could be repaired by resident or circulating stem cells, repairing massive infarcts may not be repaired with the endogenous repair processes. This leads to irreversible damage in cardiac muscles mainly via loss of cardiomyocytes, endothelial cells, and smooth muscle cells [2]. Therefore, there is an urgent need for a novel therapeutic approach to ameliorate the clinical outcome to enhance cardiac function by replacing the injured heart cells. Stem cell-based therapies have demonstrated the potential to regenerate damaged cardiac tissue directly through cell replacement or indirectly through local paracrine effects [3–5]. The survival rate and retention capacity of these transplanted cells are important parameters to optimise for a long-term therapeutic effect in a damaged heart. In addition, the post-infarction region also has an ischemic and inflammatory environment. This environment remains a notable challenge against the alterations by the transplanted stem cells, such as their adhesion, survival, differentiation into cardiac cells, and establishing connections with other cells [6, 7]. Therefore, bringing new methods that increase the efficiency of stem cell transplantation into clinical applications is highly essential.

Adult cardiac progenitor cells (CPCs) have been proposed as a potential treatment for myocardial regeneration through their high cardiomyogenic potential [8–10]. They can differentiate into cardiac cell types, such as smooth muscle, endothelium, and cardiomyocytes [11]. It is suggested that CPCs promote angiogenesis and blood flow while suppressing inflammation, fibrosis, and apoptosis in the injured tissue [12, 13]. These properties of CPCs make them appealing candidates for treating acute myocardial infarction. CPCs are collected from a cell population migrating from heart tissue explants within a few days. This population spontaneously form spherical aggregates, called cardiospheres, and is highly enriched with stem and progenitor cells, and thereby exhibits cardiovascular differentiation potential and regenerative activity [10, 14–19] These cells are also referred to as cardiosphere-derived progenitor cells. CPCs are isolated, expanded and applied by this technique in Phase 1 and 2 clinical trials [20–23].

Previous studies have demonstrated that pretreatment with cytokines, exosomes or chemical factors enhances the therapeutic efficacy of transplanted cells by modulating their survival rate, adhesion, differentiation and angiogenic abilities [24, 25] and thereby contributing to improved myocardial repair [26]. Although estrogen is not amongst these factors, it is well-known that estrogen has regenerative and cardioprotective effects on cardiac regeneration [27–29]Both estrogen receptor α (ERα) and estrogen receptor β (ERβ) are expressed in adult and neonatal hearts, although ERα is increased in cardiomyocytes after cardiac injury [30, 31]. Furthermore, ERα expression also increases in c-Kit+ CPC following myocardial infarction (MI) and stimulates proliferation of undifferentiated myoblasts [31]. In addition to cardioprotective effects, estrogen receptors have an important impact on the regulation of electrophysiological and contractile activities of the heart by controlling the expression of genes involved in ion channels and other factors contributing to excitation-contraction coupling [32]. There is a gender-dependent differences in the risk of MI. The prevalence of MI is higher for men compared to women [28]. It was also revealed that gender differences mediate the differential epigenetic regulation in endothelial progenitor cells regarding improving their myocardial ischemic repair mechanisms [33].

Recent studies also emphasized the role of epigenetic regulation in cardiovascular disease by considering its advances in clinical trials [34]. The involvement of several epigenetic mechanisms, such as DNA methylation, histone modification, and noncoding RNA expression, has been documented for cardiovascular disease development and regression. In addition, it has also been shown that estrogen signalling provides a bridge between the nucleus and mitochondria in cardiovascular diseases [35]. Finally, a recent study provided an important highlight on the importance of the sex of donor cells for progenitor-based tissue repair through the process related to the regulation of epigenetic mechanisms in sex differences associated cardiac reparative functions of bone marrow progenitor cells [33]

Considering the active roles of estrogen in cardiac remodelling, not only functional levels but also genetic and epigenetic levels, in this study, we aimed to investigate the regenerative responses of estrogen-pretreated-transplanted CPCs both in the mouse heart failure model and in vitro. Our results provide that estrogen pretreatment induced cardiac recovery by (1) promoting engraftment (2) increasing proliferation, differentiation and angiogenesis (3) attenuating morphological and structural cardiac remodeling (4) maintaining higher mitochondrial energy (5) regulating gene expression involved in cell cycle, DNA repair, glycolysis, mitochondrial biogenesis, epithelial-mesenchymal transition (EMT) and ROS production and immune response and (6) improving of cardiac functions as a result of the regulation of all these mechanisms. The current study suggests that estrogen pretreatment may help overcome the low regenerative efficacy observed in autologous/allogeneic transplantation with CPCs, which are also used in clinical trials.

## Methods

### Cardiac progenitor cell isolation from mice and culture

CPCs were cultured from the hearts of 4-6-weeks old BALB/c male mice as described, previously ([10, 15, 19, 36]). Briefly, hearts were collected in cold DMEM/F-12 containing 2% Penicillin-Streptomycin (P/S) and washed multiple times in cold Dulbecco’s PBS (Gibco, 14190-144) containing 4% P/S. The tissue was cut into 2-3 mm^3^ fragments and then partially digested in pre-warmed DMEM/F-12 containing 0.1% collagenase IV (Sigma, C5138) and 0.0001% trypsin-EDTA at 37°C for 20 min. After multiple washes in explant medium (IMDM supplemented with 0.1 mmol 2-mercaptoethanol (Gibco, 21985023), 20% FBS, 100 U/ml penicillin, 100 μg/mL streptomycin and 2 mmol/L L-glutamine), these minced cardiac explants were placed on 6-well plates precoated with fibronectin (20 μg/ml) in explant medium. Next day, 1.5 mL explant medium was added dropwise on top of each well. In 7-10 days after tissue explanting, cardiac outgrowths surrounding the explant were harvested by incubating the cells with 0.1% trypsin-EDTA at 37 °C for 7 min and then were seeded on fibronectin-coated dishes in CPC growth medium (DMEM/F-12 including 20 ng/ml bFGF (BIOSOURCE,PHG0266), 20 ng/mL EGF (BIOSOURCE,PHG0314), 10 ng/mL Hlif (BIOSOURCE,PHC9484), 2% B27 (Gibco, 17504001), 10% FBS, 100 U/mL P/S, 2 mmol/L L-glutamine). Cardiac outgrowth could be harvested up to 5 times from the same specimen. CPCs pretreated with 10^-7^ M estrogen (Sigma, E2758,) four 48 hours (h)[37, 38]

### Heart failure mouse model

Heart failure model was generated by isoproterenol (ISO) injection into BALB/c female mice (19±3 g, 6-8 weeks). 200 mg/kg ISO injected as inraperitionally daily for 6 consecutive days) [39]. Femal mice group were divided into four groups: Sham group (saline-injected), isoproterenol group (ISO) (isoproterenol induced-heart failure group), +Control-CPC group (untreated-CPCs transplanted group following ISO administration); +E2-CPC group (estrogen-treated CPCs transplanted group following ISO administration). On the day after the last ISO administration, all animals under wun treatedent baseline electrocardiogram (ECG) recordings using a PL3504 PowerLab 4/35 data acquisition system (AD Instruments, New South Wales, Australia) to confirm heart damage. Intramyocardial transplantation was performed through ultrasonography.

### Labeling CPCs with Dil and intramyocardial injection

To follow the engraftment of the cells after transplantation, the cells stained with fluorescence-labeled Dil dye just before injection. Estrogen-treated CPCs (E2-CPCs) and untreated-CPCs (Control-CPCs) were labeled by CM-DiI (Invitrogen, C7001). A 10 μL of the dye solution dissolved in DMSO (1 mg/mL) was added into 1 x 10^6^ cells/1 mL PBS, the cells were incubated at 37°C for 30 minutes (min). Following the washing step 1 x 10^6^ cells/group were dissolved in 100 μL saline to intramyocardial injection. Intramyocardial injection previously reported as a route, which provided long-term survival, retention and benefits of transplanted cells [40–42]. The injected cells were visualized with TRITC filters 2 h and 48 h after transplantation using an imaging system (IVIS, Perkin Elmer) in the live animal system.

### Fluorescence in situ hybridization (FISH) – Immunfluoresence combined staining

FISH combined with immunohistochemistry for tissue-specific markers provides a reliable method for monitoring the fate of somatic stem cells in transplantation experiments [43]. Serial tissue sections taken from frozen heart tissue blocks were blocked with Alexa Flour 594 fluorescently labelled Y chromosome probe (Cambio, CT-WPP124) mixture and coverslips were closed with rubber cement. Slides were heated at 77°C for 3 min on a heating plate and incubated overnight at 37°C in a humidified container. After the coverslip and adhesive were removed, the slide was first washed in 0.4 x Saline-sodium citrate buffer (SSC) (pH 7.0) for 2 min at 73°C, and then in 2 x SSC, 0.05% Tween-20 (pH 7.0) for 30 s at room temperature (RT). Sections marked with a PAP-pen were placed in a moist container and blocked with 5% BSA/0.025% Triton X-100/1X PBS for 1-1.5 h at RT. After draining the blocking solution on a napkin, the sections were incubated overnight at +4°C with mouse anti-cardiac troponin-I (cTntI) (Santa Cruz, sc-133117) antibody. After washing twice in 1x PBS for 5 min each, incubated with fuorescein goat anti-mouse IgG (Rockland, 610-1202) secondary antibody for 2 h at RT. Coverslips were covered with DAPI mounting media. Images were taken using an LSM-880 confocal microscope (Carl Zeiss, Oberkochen, Germany). Sections from untransplanted female mice with heart failure were used as negative controls (ISO) for Y-chromosome staining.

### Electrocardiographic and Echocardiographic analysis

Electrical activity of the heart *in situ* assessed by surface electrocardiogramme (ECG) recordings and data were acquired by using an analog-to-digital converter BIOPAC MP35 (Goleta, California) and processed with a high-cut (low-pass) filter at 50–500 Hz. Experimental animals underwent ether inhalation anesthesia during the recordings and then bipolar limb leads (lead I, II, III) were carried with carefully placed 20 gauge needles to forearms and hind limb. ECG recordings were taken for 10 min from every animal. Echocardiography was performed on anesthetized mice by 4% isoflurane using micro–computed tomography (Micro-CT) (Perkin Elmer, USA) with cardiac gating mode. Left ventricular pressure-volume loop measurements were recorded and left ventricular ejection fraction (LVEF), cardiac output (CO) and stroke volume (SV) parameters were calculated by previously decribed calculations [44]). Electrocardiographic and echocardiographic measurements were performed three different time points; before the ISO administration, at the first 1 day following ISO administration and at day 10 post-transplantation.

### Histological analysis

Mice were sacrificed at day 10 post-transplantation. Half of the heart tissue samples were fixed overnight in 3.7% paraformaldehyde for overnight and then placed in 1.2 M sucrose+ 0.1% PFA solution in PBS for 6-12 h. All samples were OCT (Tissue-Tek, 4583)-embedded and sectioned at 5 µm. For Picro-Sirius red staining, frozen sections were fixed in 70% EtOH and stained with 0.1% Sirius Red/Direct Red 80 (Sigma, 365548) solution prepared in aqueous saturated picric acid for 1 h after washing with distilled water. Sections were then washed twice with 0.5% acetic acid for 30 s/each, dehydrated three times with 100% ethanol for 1 min/each, cleared with xylene, and mounted with xylene-based mounting medium. The percentage of Sirius Red-stained area was measured by ImageJ software with an adjusted unchanged threshold.

### Transwell migration assay

A total of 1 × 10^5^ CPCs in 500 μL medium were seeded into the upper chambers of 24-well transwell inserts (8 μM pore-sized, Millipore, Billerica, MA, USA) and then incubated for 24 h at 37 °C in a humidified atmosphere with 5% CO2. After 24 h, the membranes were removed, and its upper surface was wiped away with a cotton swab to remove the not migrated CPCs. The membrane was then fixed in neutral formalin for 10 min at RT and then stained with 0.1% crystal violet for 15 min. The number of CSCs that have migrated to the lower surface of the membrane was counted under a light microscope (Carl Zeiss, Oberkochen, Germany). Each assay was performed in triplicate wells.

### In vitro angiogenesis assay

To assess the angiogenic potency of estrogen on CPCs, CPCs were seeded at 1.0 × 10^4^ cells/cm^2^ in a matrigel-coated 96-well plate (BD Biosciences, CA, USA). CPCs were incubated with 10^-7^ M estrogen for 12 h to allow the formation of tube-like structures. The cells were viewed under a light microscope (Carl Zeiss, Oberkochen, Germany); the images of the capillary network were acquired, and total tube lengths formed were measured using the ImageJ software (National Institutes of Health, MD, USA). Tube formation assays were performed in triplicate wells [45].

### Immunofluorescence assay

For immunofluorescence, sections were rehydrated with PBS for 5 min, permeabilized with 0.25% Triton X-100/PBS and blocked in 5% BSA+0.25% Triton X-100 in PBS for 1 h at room RT. Primary antibodies were diluted at 1:50 for mouse anti-Von Willebrand factor (VWF) (Santa Cruz, sc-365712), 1:200 for rabbit anti-CXCR4 (Proteintech, 11073-2-AP), 1:50 mouse anti-cTntI (Santa Cruz, sc-133117), 1:200 rabbit anti-Ki67 (Elabsience, E-AB-60601), 1:200 for mouse anti-alpha-SMA (αSMA) (Santa Cruz, sc-53015). After incubation with the primary antibodies overnight, sections were rinsed twice and incubated with Texas red goat anti-rabbit IgG (H&L) (1:500) (Rockland, 611-1902) and/or fluorescein goat anti-mouse IgG (H&L) (1:500) (Rockland, 610-1202) for 1 h at RT. Mounting Medium with DAPI (Novus Biologicals, H-1200-NB) was used for mounting and images were taken using a LSM 880 confocal microscope (Carl Zeiss, Oberkochen, Germany).

### Quantitative real-time Polymerase Chain Reaction (qRT-PCR)

Heart tissues as well as estrogen-treated or control CPCs underwent RNA extraction with PureZOL™ trizol (Bio-Rad, 7326880) and 1 µg of purified RNA was reverse-transcribed into cDNA using the iScript cDNA Synthesis Kit (Bio-Rad, 1708891). qRT-PCR was performed using the LightCycler 480 II (Roche, Basel, Switzerland) together with the SsoAdvanced^TM^ Universal SYBR Green Supermix (Bio-Rad, 1725271) according to the manufacturer’s instructions. The following primer sequences (SupplementaryTable S1) were used. Samples were amplified in duplicates and the relative gene expression levels were determined with normalization to GAPDH gene expression levels followed by the 2^−ΔΔCt^ method.

### Colony formation assay

Spheroid formation was created using AggreWell800 (Stem Cell Technologies, Canada) plates. AggreWell anti-adherence rinsing solution was added to plates and centrifuged to remove bubbles. After washing the wells, CPCs were seeded at 3×10^6^ per well with CPC media. One day later spheroid formations were visualized under the light microscope (Carl Zeiss, Oberkochen, Germany).

### Cellular membrane characterization

CPCs were labelled with 100 nM of the ratiometric mitochondrial membrane dye; Nile Red Mito (NRMito) and performed spectral imaging as described previously [46]. NRMito exhibits a strong spectral shift toward the red between ordered and disordered phases. GP values are ranges between +1 and −1 and this value is inversely proportional to membrane fluidity. Briefly, higher GP indicates lower fluidity correalated with lipid order, increased lipid packing and reduced membrane polarity. Laser light at 488 nm will be selected for fluorescence excitation of NRMito. The lambda detection range was set between 490 and 694 nm and the wavelength interval of each step has been set to 8.8 nm. The GP values were calculated as explained previoulsy [46]. We defined λ Lipid order (Lo) and λ Lipid disorder (Ld) as 570 and 650 nm for NRMito probes.

### Transcriptome analysis

RNA-Seq data analysis was initiated from raw count matrices, with DESeq2 (v1.46.0) used to normalize the data and conduct differential expression analysis. DESeq2, available through Bioconductor, applies a negative binomial distribution model, adjusting for sample-level variables such as sequencing depth and library size to ensure accurate normalization and comparability across samples. Following normalization, pathway enrichment analysis was conducted using ReactomePA (v1.50.0), which maps differentially expressed genes (DEGs) to pathways in the Reactome database. ReactomePA allows for the detection of pathways with significant enrichment in DEGs, thus highlighting biological processes potentially impacted by experimental conditions. In parallel, Gene Set Enrichment Analysis (GSEA) was performed with clusterProfiler and fgsea (v1.0). These R packages rank genes based on differential expression to identify gene sets with coordinated expression shifts across experimental conditions. For GSEA, the Molecular Signatures Database (MSigDB) database was used, with significance assessed at a false discovery rate (FDR) threshold of 0.05 to pinpoint biologically meaningful gene sets. All data visualizations were created in Python, employing several specialized libraries to ensure clarity and depth in data interpretation. Matplotlib (v3.7.1) was used for foundational plotting such as histograms and scatter plots, which illustrate broad distribution patterns within the normalized count data. For more detailed statistical visualizations, Seaborn (v0.12.1) facilitated the creation of heatmaps, boxplots, and violin plots, enriching the interpretation of differential expression results. Additionally, Plotly (v5.14.1) enabled the production of interactive visualizations, allowing an in-depth exploration of gene-specific and pathway-specific expression patterns.

### Statistics

Statistical analyses were performed using Prism 5.0 (GraphPad Software Inc.). For most experiments, three biologically independent replicates were performed, and otherwise stated differently in the corresponding figure legend. The statistical difference between the two groups was analyzed using an unpaired Student’s t-test. Differences with two-tailed p-values < 0.05 were considered statistically significant. All results are expressed as mean ± standard deviation (SD).

## Results

### Estrogen pretreatment enhances the retention rate of transplanted CPCs into failing hearts

Since a high retention rate is a prerequisite for the success of cell-based heart regeneration, we evaluated the survival and engraftment of CPCs following intramyocardial transplantation. For that purpose, either male CPCs (1×10^6^ cells) or vehicle (saline) were injected into adult female mice with heart failure. The most important reason for using male mouse cells is to observe the estrogen effect more clearly. Another reason is to observe the retention rate by monitoring the presence of the Y chromosome in the female heart. Isolated CPCs from male mice were characterized by flow-cytometry analysis (Supplementary Fig. 1A) and assessed for their ability of colony formation which reflects their self-renewal capacity (SupplementaryFig. 1B). Characterized CPCs were incubated with estrogen for 48 h and then labelled with Dil staining to detect short-term engraftment via fluorescence imaging before being injected intramyocardially into female mice with ISO-induced heart failure. The mice with heart failure (ISO) were injected with either vehicle or estrogen estrogen-pretreated CPCs (E2-CPCs) or untreated CPCs (Control-CPCs). The expression of both ERα and ERβ was verified in all groups (SupplementaryFig. 1C). Following the transplantation, the labelled cells were visualized after 2 h and 48 h. Both Control-CPCs and E2-CPCs showed a high fluorescence signal at 2 h. While control-CPCs were completely invisible at the end of 48 h, fluorescence signal intensity of transplanted E2-CPCs showed only around 34 % reduction compared to the 2-hour time point (Fig.1A). To assess long-term engraftment of transplanted male CPCs in the female heart, Y chromosome detection was perfomed through the FISH-IF combined staining [43]. The mice were sacrificed at day 10 post-transplantation. Heart tissue sections were co-stained with cTntI (green) to detect the cardiomyocytes and Y-chromosome (red) to confirm the engraftment of the male donor cells. Rare Y-chromosome positive nuclei (DAPI) were observed in Control-CPC injected hearts, while a 3-fold higher number of Y-chromosome positive cells were detected in the E2-CPC-transplanted hearts compared to control CPC transplanted recipients (Fig. 1B). Y-chromosome signal could not be detected in cTntI-positive cells in the recipient hearts of both groups, suggesting that the 10-day post-transplantation period may have been insufficient for CPCs to differentiate into cardiomyocytes. The higher retention rate of Y chromosome-positive cells was also confirmed by examination of SRY gene expression levels in transplanted female heart tissue (Fig. 1C). These results indicated that the estrogen-pretreated pretreated-CPCs demonstrated a significant increase in engraftment in failing hearts in vivo as compared to untreated CPCs.

**Figure 1.**
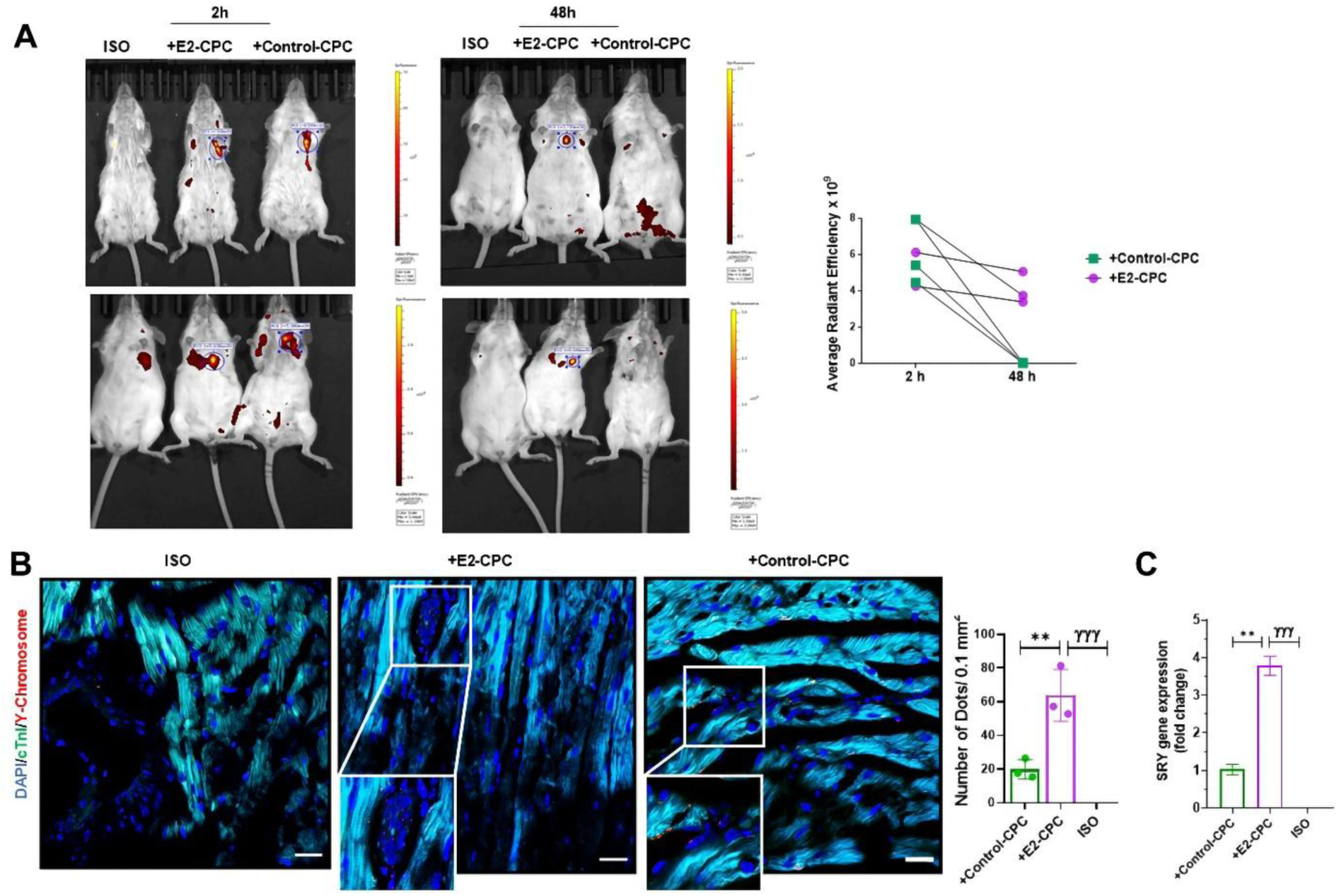
Engraftment of CPCs in the failing mouse heart. **(A)** In vivo fluorescence images of mice with ISO-induced heart failure at 2 h and 48 h after intramyocardial injection of untreated CPC (+Control-CPC group) and E2-treated CPC (+E2-CPC group). The ISO group represents the mice with ISO-induced heart failure. Vehicle (saline) was only injected into ISO group as a negative control **(B)** Detection of the Y-chromosome via FISH-IF analysis. Images are taken with 20x and 40x objectives. The white scale bars represent 20 µM. **(C)** mRNA levels of the SRY gene by qRT-PCR analysis in CPC transplanted female mouse hearts. n=3 (biological replicates). Data are presented as mean±SD. **P < 0.001 *vs* +Control-CPC, ^ᵞᵞᵞ^P < 0.0001 *vs* ISO.

### E2-CPC transplantation improves cardiac function in a failing heart

To demonstrate the cardiac electrical functions, echocardiographic and electrocardiographic analyses were used to assess the functional effects of transplantation of E2-pretreated CPCs. All functional analyses were performed on day 10 post-transplantation. ISO-induced heart failure has also been confirmed by expression of heart failure and fibrosis markers (Suppl. Fig 1D), in addition to echocardiographic and electrocardiographic evaluations. ISO administration profoundly reduced LVEF, SV, and CO, indicating LV dysfunction in failing female hearts. E2-CPC treatment resulted in by 2-fold increase in LV function. Conversely, Control-CPC injection could not reverse the LV dysfunction caused by ISO administration (Fig.2A). Heart failure conditions such as ischemia and atrial fibrillation are characterized by progressive and intrinsic decline in cardiac function, typically reflected by prolonged QT and PR intervals on surface electrocardiograms (ECGs), reduced maximum heart rate and reduced contractile activity as previously shown [47–49]. The durations of PR, QRS, and P-wave were detected as significantly decreased in both control CPC and E2-CPC transplanted hearts. E2-CPC injection signficantly reduced QT and PR intervals compared to control CPC injected groups (Fig. 2B). Consistent with the echocardiographic and electrocardiographic findings, heart weight to body weight ratio (HW/BW) used as an index of cardiac hypertrophy, was also observed as dramatically reduced after E2-CPC engraftment (Fig. 2C).

**Figure 2.**
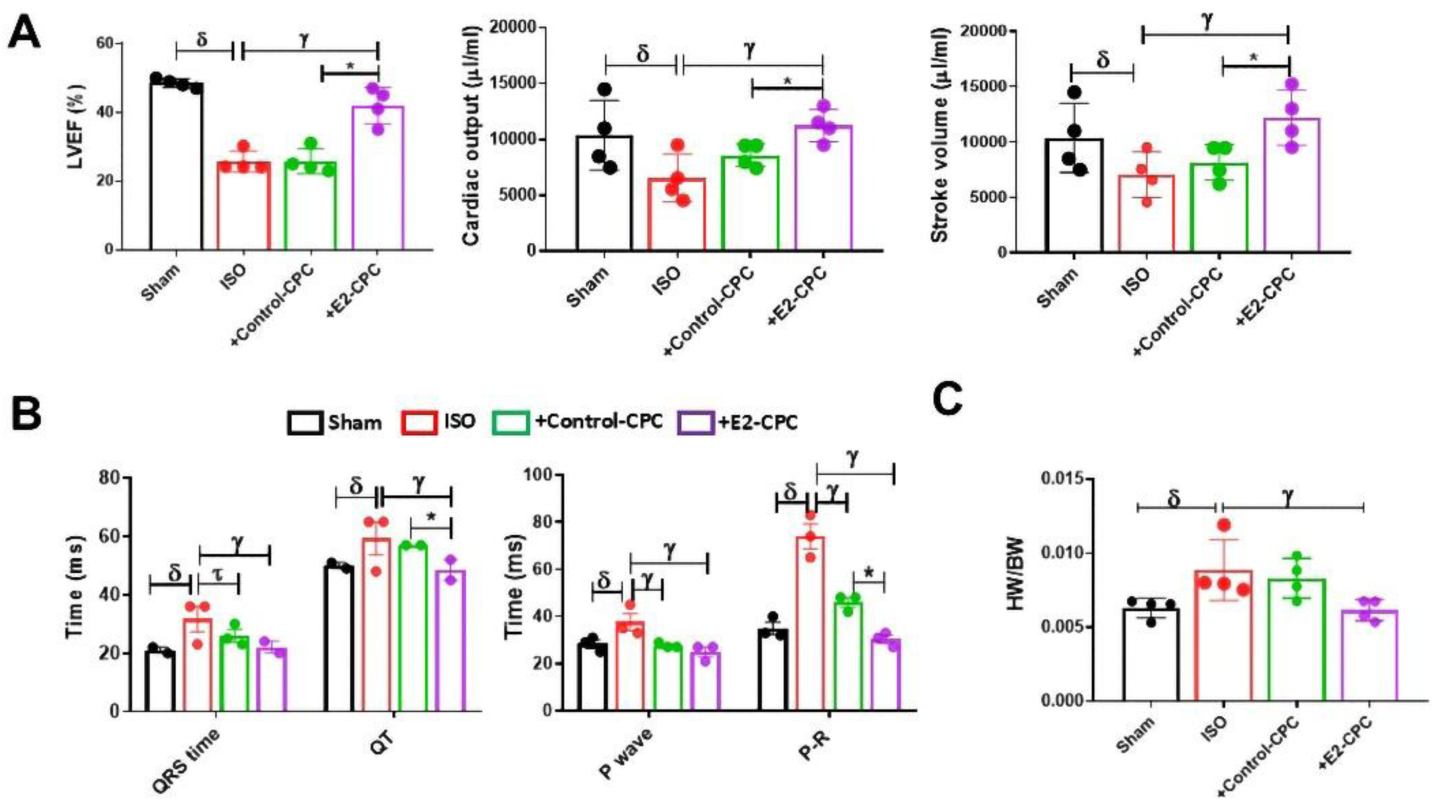
Evaluation of cardiac functions following the CPC transplantation. **(A)** Echocardiographic analysis of either in E2-PC injected group or the Control-CPC injected group. Sham represents the only vehicle-injected healthy control group. The echocardiographic analysis was performed by Micro-CT n=4(biological replicates). **(B)** Bar graph of PR, QT, QRS intervals and P waves recorded by ECG analysis and **(C)** The HW/BW ratio of SHAM, ISO, +E2-CPC and +Control-CPC mice. n=3 (biological replicates). Data are presented as mean±SD.*P < 0.05 *vs* +Control-CPC, ^ᵞ^P < 0.05 *vs* ISO, ^δ^P < 0.05 *vs* Sham.

### E2-CPC transplantation stimulates cardiac recovery, promoting revascularisation and proliferation while attenuating collagen accumulation and fibrosis

Cardiac remodelling in heart failure serves as an important compensatory mechanism of congestive heart failure, characterised by progressive hypertrophy, fibrosis, loss of functional cardiomyocytes and reduced neoangiogenesis [50]. Here, we assessed the revascularisation in failing hearts by Von Willebrand factor (VwF,) and cardiomyocyte proliferation was evaluated by cardiac troponin I (cTnI) expression level. Although ISO administration caused in significant reduction in both VwF and cTnI expression compared to the Sham group, whereas E2-CPC transplantation markedly enhanced the protein levels of these two parameters. Importantly, no significant differences were observed between Control CPC transplanted and ISO hearts (Fig. 3A). As shown in Fig. 3A, alpha-smooth muscle actin (α-SMA) fibre expression decreases in E2-CPC transplanted hearts compared to Control CPC transplanted hearts. Furthermore, we could not detect CXCR4 expression in both ISO and CPC-injected groups, but drastically high expression was observed in E2-CPC engrafted heart tissue, especially in the atrium. We also evaluated the cell-cycle activity in the heart by determining Ki67. ISO administration drastically abolished the Ki67 expression, while CPC transplantation could slightly reverse this impaired proliferation. Furthermore, there was the highest expression of Ki67+ in E2-CPC transplanted hearts compared to the other groups (Fig. 3B), indicating the induction of cell proliferation by E2-CPC transplantation. Considering the role of nuclear factor of activated T cells 3 (NFATc3) on the development of hypertrophy and collagen type I alpha 1 chain (Col1a1), we determined the protein level of NFATc3 in transplanted heart tissues. Although CPC transplantation reduced NFATc3 and Col1a1 protein levels in recipient hearts, E2-CPC caused a more significant reduction in NFATc3 and Col1a1 protein levels compared to those of controls (Fig. 3C). Given the prevalence of myocardial fibrosis and abnormalities in collagen content of the myocardium across various cardiac hypertrophy models, here, an existence of the extensive collagen deposition characterized fibrosis is determined by Sirius red staining. Our results indicated a higher portion of collagen deposition in the ISO groups. Although CPC injection caused a slight but not significant reduction, E2-CPC injection resulted in a significant decrease in collagen deposition (Fig. 3D). These findings provide additional evidence that E2-CPCs attenuate the cardiac remodelling both at the physiological and pathological levels.

**Figure 3.**
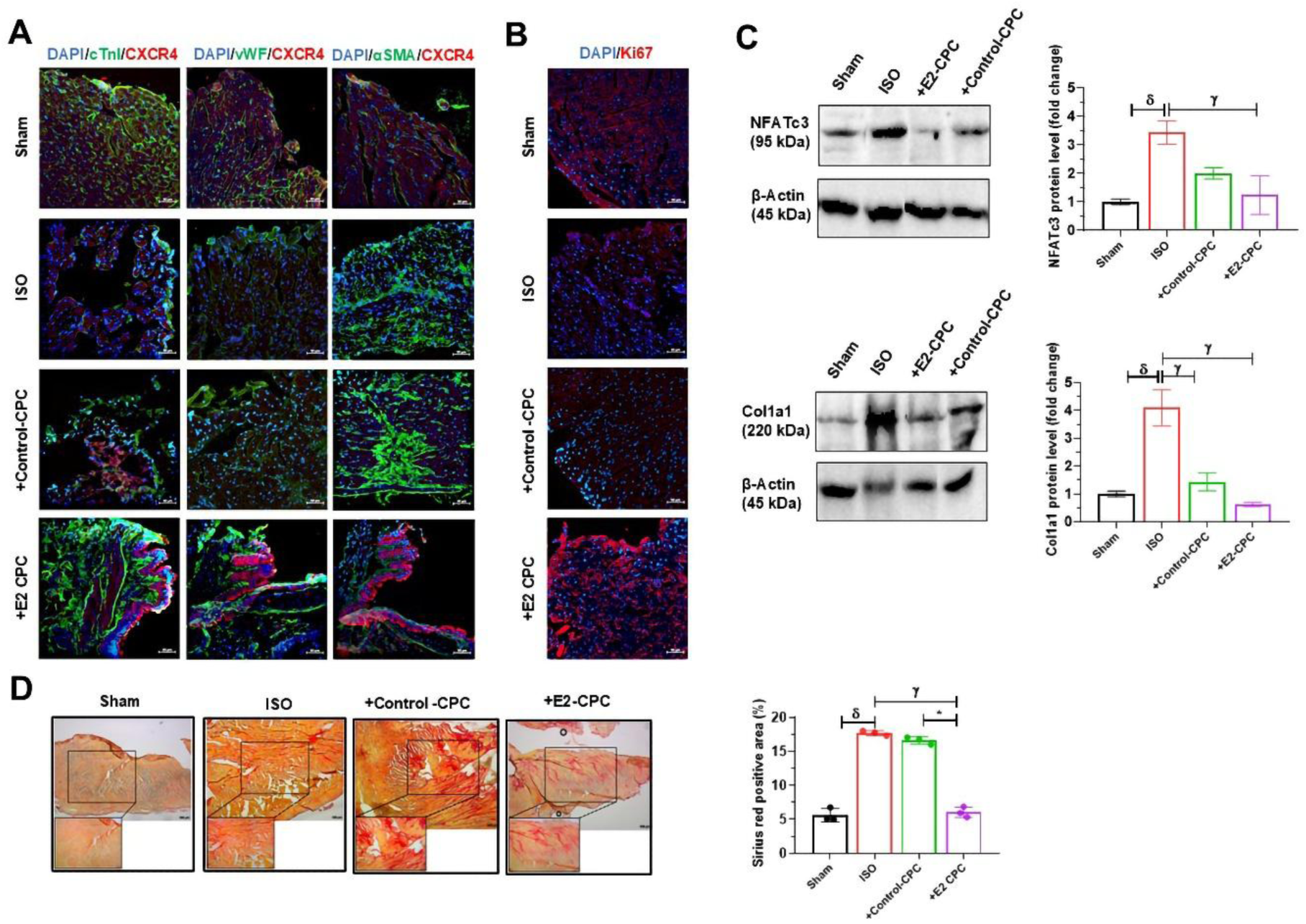
Morphological analysis of heart tissues following the CPC transplantation. **(A)**Representative immunofluorescent staining of cardiac cell markers: cTnI, vWF, αSMA, CXCR4 and **(B)** proliferation marker Ki67 on heart tissue sections. Images are taken with a 20x objective. Scale bars represent 50 µM. **(C)** Western blotting for hypertrophy and fibrosis markers, NFATc3 and Col1a1. β-Actin was used as a loading control. **(D)** Representative Sirius red staining on heart tissue sections. Images: 20x and 40x. Scale bars represent 200 µM. n=3 (biological replicates). Data are presented as mean±SD.*P < 0.05 *vs* +Control-CPC, ^ᵞ^P < 0.05 *vs* ISO, ^δ^P < 0.05 *vs* Sham.

### Estrogen treatment induces migration, angiogenesis and mitochondrial energy of CPCs *in vitro*

Following the detection of high potentiality of estrogen pretreatment as a possible therapeutic agent, we focused on understanding the molecular and cellular underpinnings of the changes in CPCs upon estrogen treatment. For this aim, we first investigated the effect of estrogen treatment on the migration capacity of CPCs through the transwell membranes. Estrogen application highly stimulated the migration ability of CPCs (Fig. 4A). In addition to the control and E2-CPC groups, we also analyzed the migration capacity of CPCs, which were isolated from male mice with heart failure induced with ISO (ISO-CPC). Among these groups, a few numbers of migrated ISO-derived CPCs were detected. The enhancer role of estrogen in the migration of CPCs was also demonstrated by wound healing assay (SupplementaryFig.2). In addition to migration capacities, the tube formation ability of CPCs was also investigated to assess their angiogenic potential. Our results showed a significant recovery in the angiogenic capacity of CPCs following estrogen treatment (Fig. 4B). It is known that the interaction of SDF-1 and CXCR4 enhances the expression of angiogenic and migration signals [51]. Furthermore, increased SDF1 and CXCR4 expression levels in these samples correlate an enhancement in the angiogenesis and migration capacities of CPCs under estrogen treatment (Fig.4C). Considering estrogen treatment associated with an increase in genes involved in proliferation including ABCG2, Ki67, and Nkx2.5, we further aimed to evaluate the regulatory function of estrogen on the differentiation capacity of CPCs. To this end, we analyzed the expression of cardiomyocyte and endothelial differentiation markers; cTnT and vWF, respectively. The estrogen treatment increases the expression of both cTnT and vWF, which suggests that estrogen may play a role in the differentiation of CPCs into cardiac cell types (Fig.4C).

**Figure 4.**
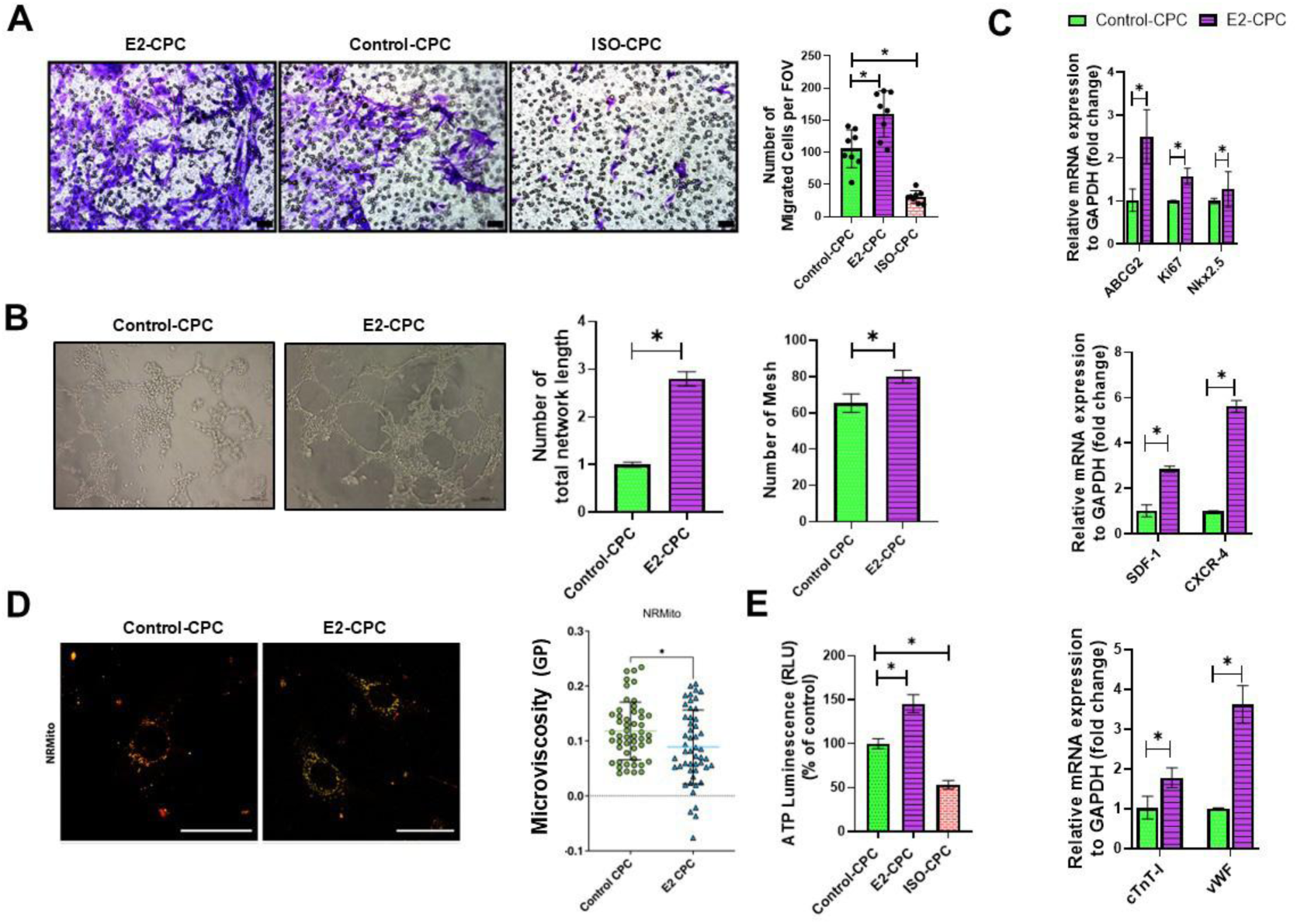
In vitro analysis the effects of estrogen treatment in CPCs. **(A)** Transwell assay showing migration of CPCs in vitro. Cells were stained with crystal violet. ISO-CPC represents isolated CPCs from ISO-induced hearts. Images are taken with 20x objective. The black scale bars represent 100 µM. **(B)** Representative images of the tube formation assay. Tube length and number of mesh are calculated and represented with bar graphs. Images are taken by 20x. The black scale bars represent 100 µM. **(C)** mRNA levels of SDF1, CXCR4, ABCG2, Ki67, Nkx2.5, cTnI and vWF, compared to GAPDH by qRT-PCR analysis. **(D)** GP values of mitochondrial membranes. Images are taken by 20x.The white scale bars represent 100 µM. Images are representative of technical replicates **(E)** ATP production in CPCs. (n=3)(biological replicates). Data are presented as mean ± SD. *P < 0.05 *vs* Control-CPC.

Mitochondrial dynamics is also crucial for stem cell homeostasis, pluripotency, differentiation and cell fate decision, consequently [52]. To this end, we have also examined the effect of estrogen treatment on the mitochondrial energetics. Estrogen induces ATP production in CPCs (Fig. 4D). In advance, mitochondrial membrane fluidity is also a crucial aspect of mitochondrial energetics, as it influences the functionality of the electron transport chain and the efficiency of ATP synthesis. For this purpose, mitochondrial membranes were labelled with NRMito ([53, 54] and GP values were recorded as described before ([46]). Our result has shown that estrogen treatment reduced the lipid order in the mitochondrial membrane of CPC (Fig. 4D). These results suggested that changes in mitochondrial dynamics could be the other important factor which enhances the higher regenerative capacity of transplanted E2-CPCs.

### Estrogen changes the transcriptome profile of CPCs

To explore the key genes and pathways related to cell survival and retention, cardiac remodelling, migration, differentiation and mitochondrial function, differentially expressed genes (DEGs) between control and estrogen-treated CPCs were examined. The biological roles of common DEGs were explored through enrichment analysis. The overall workflow of this study is shown in Figure 5. DEGs were identified following normalization of RNA-seq data. Significance analysis of the RNA-Seq results revealed a total of 675 genes that were differentially expressed in estrogen-treated CPCs (≥1.2-fold, P<0.01); 272 genes were upregulated and 403 genes were downregulated, as represented volcano plot (Fig. 5A). The most signifcant estrogen regulated up-regulated DEGs were involved in response to supression of oxidative stress and mitochondrial dysfunction (e.g., *SOD3*, *Isg15*), inhibition of myocardial apoptosis (cFOS), cell adhesion and retention (Icam1), attenuation of the immune response and supression of cardiac dysfunction and hypertrophy (Usp18). Heatmap provides an overview of the top 1,000 DEGs also demonstrating differentially expressed gene sets between control and estrogen-treated CPC groups (SupplementaryFig. 3).

**Figure 5:**
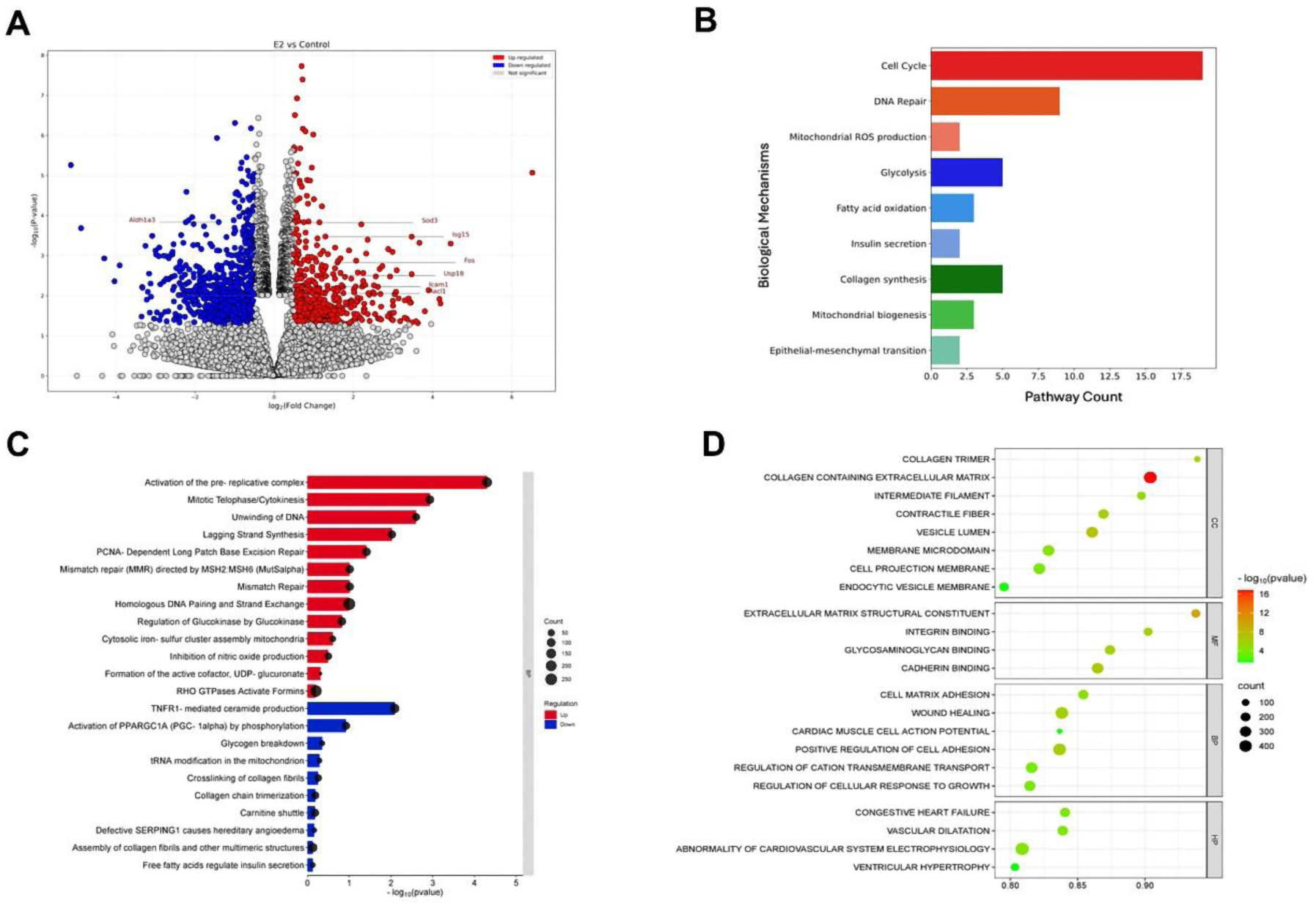
RNA-seq of E2-CPC vs Control-CPC groups. **(A)** Volcano plot showing differentially expressed genes between E2-treated and Control groups, highlighting significantly upregulated (red) and downregulated (blue) genes. **(B)** Summary of related pathways affected by E2 treatment, showing the count of genes involved in key biological mechanisms. **(C)** Reactome pathway analysis showing significantly enriched pathways in E2-treated vs Control groups. The size of each black circle indicates the number of genes corresponding to each pathway. **(D)** A bubble plot depicts GSEA enrichment analysis. The horizontal axis of the bubble plot represents GeneRatio in differential expression. The vertical axis represents the enriched pathway name. The size of each bubble indicates the number of genes corresponding to each pathway. The colour of the bubbles represents the significance of the P-value, with darker colours indicating a smaller-log10(P-value) and thus a more significant enrichment.n=3 (biological replicates)

Since estrogen modulates the gene expression profile in CPCs, we hypothesised that examining the estrogen-regulated DEGs would improve our understanding of the induced regenerative capacity of transplanted CPCs depending on estrogen treatment. To gain insight into the functional significance of estrogen-regulated up-and down-regulated DEGs, we conducted pathway enrichment analysis using both ReactomePA (v1.50.0) and GSEA pathway analysis. In Figure 5B, a summary of the enrichment analysis of biological pathways using ReactomePA for DEGs is represented. Estrogen-regulated DEGs were mostly related to cell cycle, DNA repair, collagen syhthesis, glycolysis, FA oxidation, mitochondrial biogenesis, epithelial-mesenchymal transition (EMT), insulin secretion and ROS production. Reactome pathway analysis demonstrated that the estrogen-regulated up-regulated DEGs were associated with cell cycle process pathways including pre-replicative complex activation, cytokinesis; DNA repair mechanisms including mismatch and homoglous recombination repair; mitochondrial function and mitochondrial respiratory chain including glucokinase, iron-sulfur cluster assembly; regulation of cell migration including Rho GTPase activated formins; and oxidative stress related pathways including inhibition of NO production. On the other hand, estrogen-regulated down-regulated DEGs were associated with mitochondrial dysfunction related pathways including ceramide production, carnitine shuttle, tRNA modifications in mitochondrion; collagen formation including collagen trimerization and crosslinking and assembly of collagen fibrils (Fig.5C). Different from ReactomePA analysis, GSEA functional enrichment analysis was performed based on cardiovascular system pathways and categorized into four branches: Cellular Component (CC), Biological Process (BP), Molecular Function (MF) and Human Phenotype (HP). The CCs, BPs and MFs analysis revealed that the DEGs were enriched mostly in collagen-containing extracellular matrix, extracellular matrix structural constituent, cadherin binding and positive regulation of cell adhesion pathways in estrogen-treated groups compared to controls. In the CC group, vesicle lumen formation-related genes were significantly enriched. BP analysis revealed that the DEGs were major enriched in wound healing, while in the HP group, DEGs were major enriched in congestive heart failure and vasodilatation. Interestingly in HP analysis also revealed that the number of DEGs which are related to cardiovascular system electrophsiology were observed as significantly higher in estrogen-treated CPCs, suggesting that estrogen may promote the differentiation of CPCs toward a cardiomyocyte lineage (Fig.5D). Emerging evidence suggests that the estrogen treatment changes the transcriptional profile and provides CPCs with the properties that allow them to adhere, proliferate, survive, and even differentiate in the damaged area after transplantation.

## Discussion

The introduction of CPCs prompted numerous researchers to explore their potential for experimental and clinical use as a therapeutic agent with a lower oncogenic risk and differentiation ability into multiple cardiac cell types [40, 55]. CPC transplantation has shown promising potential in preclinical trials for promoting tissue regeneration. However, translation into clinics has been hindered by challenges due to the low engraftment rate and poor understanding of the different mechanisms through which CPCs exert their therapeutic effects [56]. To advance using CPCs in cardiac regeneration, enhancing their therapeutic efficacy by regulating the engraftment, survival, differentiation, and migration mechanisms of the transplanted cells is required. In the current study, we investigated the effect of estrogen-pretreatment on the therapeutic potential of cardiac progenitor cells.

Despite the demonstrated potential of cell therapy in the treatment of cardiovascular diseases, particularly in myocardial infarction, one of the challenges that remains to be addressed is the poor efficacy of the transplanted cells [57–59]. Preclinical studies have shown that some of the transplanted cells are retained in the myocardium for minutes to hours, with a retention rate ranging from 1% to 20% [56]. Despite of the challenges in intramyocardial delivery, such as washing out of cells caused by mechanical damage of needle injection [60]Using intramyocardial delivery of cells result in higher cell retention compared to intracoronary or intravenous delivery. Due to this reason, increasing cell retention should be given as the first attention in cell-based transplantation. Here, we first examined the effect of estrogen-pretreatment in the engraftment rate of CPCs. Although control-CPCs completely diffused within the 48h following the injection, only 20% of the transplanted E2-CPCs were diffused. In addition, Y-chromosome tracking demonstrated that 10-day post-transplantation E2-CPC cells could also be detected as 3-fold higher than control-CPCs. The presence of Y-chromosome signal was not detected in cTntI-positive cells within the recipient hearts of both groups. This finding can be attributed to the insufficient duration of the 10 days post-transplantation, which was inadequate for the CPCs to undergo differentiation into cardiomyocytes within the recipient heart.

It is known that higher transplantation efficiency is associated with better recovery. E2-CPC treatment significantly improved characteristics of left ventricular function and contractile activity as well as reduced heart fibrosis and cardiac hypertrophy at 10 days post-transplantation, as opposed to control-CPC treatment. We confirmed this with the functional recovery using histological and morphological analysis. Abnormal collagen content is an important hallmark of maladaptive hypertrophy and cardiac fibrosis. Despite the observation of a minimal yet non-statistically significant reduction in collagen deposition in transplanted heart tissue following the administration of CPC injection, the administration of E2-CPC injection resulted in a significant decrease in collagen deposition. Activation and differentiation of fibroblasts into myofibroblasts is associated with pathological cardiac remodelling via collagen secretion [61, 62]. The reduced protein level of α-SMA in the E2-CPC transplanted heart may also be associated with the decreased collagen level. Cardiomyocytes and endothelial cells are essential for both heart remodelling and regeneration [63, 64]. E2-CPC transplantation increased revascularisation, as well as proliferation rate, in failing hearts. Limited cell proliferation is another major obstacle in cardiac remodelling. We detected that E2-CPC transplantation promoted the proliferation of the resident cells in cardiac tissue. Moreover, cardiac-specific chemokine receptor, CXCR4 is necessary for the proper development of the embryonic heart [65] It was demonstrated that knocking-out of CXCR4 promotes hypertrophy and cardiac dysfunction in response to chronic catecholamine exposure in an isoproterenol-induced (ISO) heart failure model [66]. CXCR4 expression was detected in a dramatically increased high in E2-CPC transplanted hearts compared to control-CPC injected hearts. These findings point out that E2-CPC transplantation enhances cardiac repair, supporting multi-factorial processes including promoted vascular formations and cardiomyocyte proliferation, reduced hypertrophy and fibrosis and subsequently improved cardiac functions as a functional consequence of all these.

Following the in vivo evaluation of the estrogen-pretreated CPC injection, we have also focused on the molecular and cellular mechanisms underlying cardioprotective changes in CPCs upon estrogen treatment. CPCs were isolated from healthy male mice to evaluate and compare regenerative capacities in response to estrogen in vitro. It was detected that estrogen induces the migration capacity of CPCs, as well as stimulating angiogenic response. Moreover, CPCs isolated from male mice with heart failure (ISO-CPC) demonstrated very limited migration capacity compared to CPCs from healthy mice, pointing out the low therapeutic efficacy of resident CPCs in the injured heart. It is acknowledged that the interaction of SDF-1 and CXCR4 enhances the expression of angiogenic and migration signals [48]. Our qRT-PCR results may also provide supportive data for the estrogen-induced angiogenesis and migration of CPCs, as well as the proliferation via increasing expression of ABCG2, Ki67, and Nkx2.5. Finally, the increased expression of cTnT and vWF upon estrogen treatment suggests that estrogen may contribute to the differentiation of CPCs into cardiac cell types. Mitochondrial dynamics are also crucial for stem cell homeostasis, pluripotency, differentiation and cell fate decision, consequently [52]. The fluidity of mitochondrial membranes, influenced by phospholipid composition and packing, critically affects membrane dynamics and electron transport efficiency [67, 68]. It was demonstrated that estrogen regulates mitochondrial energy homeostasis localising mitochondrial membranes [69]. Estrogen treatment decreased the lipid order and fluidity in the mitochondrial membrane of CPC. The decreased mitochondrial membrane fluidity promotes lateral movement of proteins involved in electron transport [70]. Collectively, our results indicate that estrogen induces cellular metabolism in a regenerative manner, mediating bionergetic homeostasis in CPCs.

Our RNA-seq data corroborate with the in vivo results. The DEG, which are upregulated following estrogen treatment, are associated with critical processes involved in attenuation of cardiac remodelling, such as supression of oxidative stress, mitochondrial dysfunction, myocardial apoptosis, cell adhesion, retention and suppression of cardiac dysfunction and hypertrophy. Pathway enrichment analysis supported the changes in molecular mechanisms, including proliferation, retention, migration and vesicle lumen formation, in response to estrogen. Especially, the presence of DEGs that are enriched in collagen-containing extracellular matrix, cadherin and integrin binding and cell adhesion pathways in estrogen-treated groups supports the results of long engraftment of transplanted CPCs in failing heart.

Estrogen treatment also causes changes in metabolic pathways, including glycolysis and fatty acid metabolism. Metabolic switch of stem cells from a glycolytic to an oxidative state is considered a key event both in the reprogramming/differentiation process [71, 72]and regulation of energy metabolism, especially under stress conditions [73]. Among DEGs, HACL1, which is involved in fatty acid oxidation was detected as the most significant estrogen-regulated up-regulated DEGs. In addition, ALDH1A3, which is involved in glycolysis, is significantly downregulated in estrogen-treated CPCs, suggesting that estrogen induces changes in metabolic pathway preferences from glycolysis to fatty acid oxidation. It is well known that the therapeutic potential of stem cell-based therapies allows for the regeneration of damaged cardiac tissue either directly or indirectly through paracrine effects [3, 74]. Considering these changes both at the molecular and transcriptome level of CPCs upon estrogen treatment in vivo, we suggested that transplanted E2-CPCs may provide paracrine effects to augment the survival of resident cardiac cells, revascularisation and facilitate beneficial remodelling, resulting in an overall improvement of heart function.

## Conclusion

We show that estrogen pretreatment may overcome the low regenerative efficacy of autologous/allogenic stem cell transplantation. Our data demonstrate that estrogen-pretreated CPCs can promote engraftment, cell survival, proliferation, endothelial cell activation, and cellular migration. We also show that both cellular and mitochondrial membrane lipid composition of CPCS may potentially contribute to these regenerative processes. We reveal that pretreatment of transplanted cells with estrogen induces changes in both the transcriptome profile and the membrane structure of the CPCs. These changes not only enable longer engraftment of injected CPCs into the damaged area but also stimulate cardiac recovery through direct or paracrine mechanisms.

## List of abbreviations

CPC: Cardiac progenitor cells
E2: 17β-estradiol
ISO: Isoproterenol
ERα: Estrogen receptor α
ERβ: Estrogen receptor β
GO: Gene ontology
KEGG: Kyoto encyclopedia of genes and genomes
GVAS: Gene Set Variation Analysis
DEGs: Diferentially expressed genes
FDR: False discovery rate
RNA-seq: RNA sequencing
FISH: Fluorescence in situ hybridization
qRT-PCR: Quantitative real-time polymerase chain reaction
Lin: Lineage cocktail
SSEA-1: Stage-specific embryonic antigen-1
ColA1: Collagen type I alpha 1 chain
NFATc3: Nuclear factor of activated T cells 3
SDF-1: The stromal cell-derived factor 1
CXCR4: C-X-C Motif Chemokine Receptor 4

## Availability of data and materials

Data are available under PRJNA1242725 accession number, https://www.ncbi.nlm.nih.gov/bioproject/1242725.

## Competing interests

The authors declare that they have no competing interests

## Funding

This study was supported by the Scientific and Technological Research Council of Turkey (SBAG-220S910). We also gratefully acknowledge the support of the European Molecular Biology Organization (EMBO) to support the stay of D.A. at Science for Life Laboratory, Karolinska Institutet by EMBO Short-Term Fellowship.

## Author Contributions

All authors have approved this manuscript and its contents, and they are aware of the responsibilities connected with authorship. C.V.B. contributed to the conception and design of the study and reviewed and approved the final version of all data. E.S. and Z.B.A. reviewed and approved the final version of the manuscript, and E.S. supported membrane fluidity experiments, imaging, and RNA-seq analysis.

D.A. performed the majority of the experiments and their analysis. N.B. completed most of the in vivo and in vitro experiments. K.G. contributed to the in vivo experiments; Z.B.A. and T.A.K. contributed to the in vitro experiments. Y.O. contributed to the in vivo electrophysiological analysis. O.E. performed the cell transplantation experiments. R.U. carried out bioinformatic analysis. T.S. contributed to and assisted with membrane fluidity measurement experiments.

## Ethics approval and consent to participate

For animal experiments, the study entitled “*Investigation of the Role of Estrogen on Cardiac Progenitor Cell Differentiation in vivo and in vitro*” was approved by the the local ethics committee of Ankara University (No. 2018-18.22) and all animal experimental procedures were performed by the European Community guidelines on the care.

## Acknowledgement

All animal experiments and wet-lab experiments were performed at Ankara University, Stem Cell Institute; electrocardiogram analysis was performed at Ankara University, Faculty of Medicine, Department of Biophysics. The authors especially thank the SciLifeLab, Karolinska Institutet, for their support with advanced microscopy imaging and RNA-sequencing.

## Consent for publication

No applicable.

## Other declarations

The authors declare that they have not used Artificial Intelligence in this study

## Authors and Affiliations

Ankara University, Stem Cell Institute, Ankara, Türkiye; Ankara University Faculty of Medicine Department of Biophysics, Ankara, Türkiye; Johannes Gutenberg University, Mainz, Germany; Veterinary Faculty Ankara University, Ankara, Türkiye; Science for Life Laboratory Department of Women’s and Children’s Health, Karolinska Institutet, Tomtabodavagen 23, 17165, Solna, Sweden.

## Supplementary Information

### Flow cytometry

Surface markers expressed on CPCs were analysed by flow cytometry. Harvested and passaged cells were resuspended in cold FACS staining buffer consisting of PBS supplemented with 3% BSA/0.05% NaN_3_ and aliquoted as 1 x 10^5^ cells/100 µL cell suspension in each flow tube. The following antibodies and conjugated fluorochromes were used: c-Kit-PE (Biolegend, 105808), CD45.1-FITC (Elabscience, E-AB-F1184C), CD90.2-PE (#E-AB-F1094D, Elabscience), Lineage (Lin) cocktail-FITC (Biolegend, 133301), CD31-APC (Elabscience, E-AB-F1180E), SSEA-1-BV421 (Biolegend, 125613), Sca-1-FITC (Biolegend, 108106), Nkx-2.5-FITC (Santa Cruz, sc-376565). The percentage of positive cells was defined as the percent of the population falling above the 99 percentile of an unstained cell population. Data were acquired on ACEA NovoCyte Flow Cytometer (Agilent, CA, USA). The analysis was performed with NovoExpress v. 1.3.0. (Agilent, CA, USA).

### Western blotting

The cells were gently washed two times with iced PBS and harvested at 4°C in lysis buffer containing 250 mM NaCl, 1% NP-40, and 50 mM Tris-HCl; pH 8.0 and 1XPIC. Total protein lysates were evaluated by BCA assay (Thermo Fisher, USA), and 30 µg protein from each group was loaded to 10 % SDS-PAGE gel. Following the transfer and blocking steps with 1% BSA in TBS-0.3% Tween, the membrane was incubated with the primer antibodies of either Col1a1 (Proteintech, 25870-1-AP), Nfatc3 (Thermo,PA5-78403), ERα (Santa Cruz,sc543) and ERβ (Santa Cruz,sc340243). β-Actin (Santa Cruz, sc-47778) primer antibody to use as house-keeping loading control. Anti-rabbit (Abcam, ab150077) was used as secondary antibody. Band intensities were calculated using Image J (NIH, USA). Band intensities were normalized according to β-Actin levels.

### Wound healing (scratch) assay

a total of 1×10^5^ CPCs were seeded in DMEM-F12 starvation medium with 0.5% FBS for 24 h in 6-well plates and allowed to form a monolayer overnight at 37 °C in a 5% CO2 incubator. Using a pipette tip, scratches were made in each well of the confluent monolayer. The medium was then replaced with DMEM-F12 containing 10^-7^ M estrogen. The wound closure was then monitored at 6 h, 12 h, and 24 h of incubation by means of an inverted light microscope (Zeiss AxioVert, Germany). Migration was quantified by measuring the width of the cell-free zone using the wound healing analyser plug-in for ImageJ (NIH, USA).

---

**Suppl. Figure 1.**
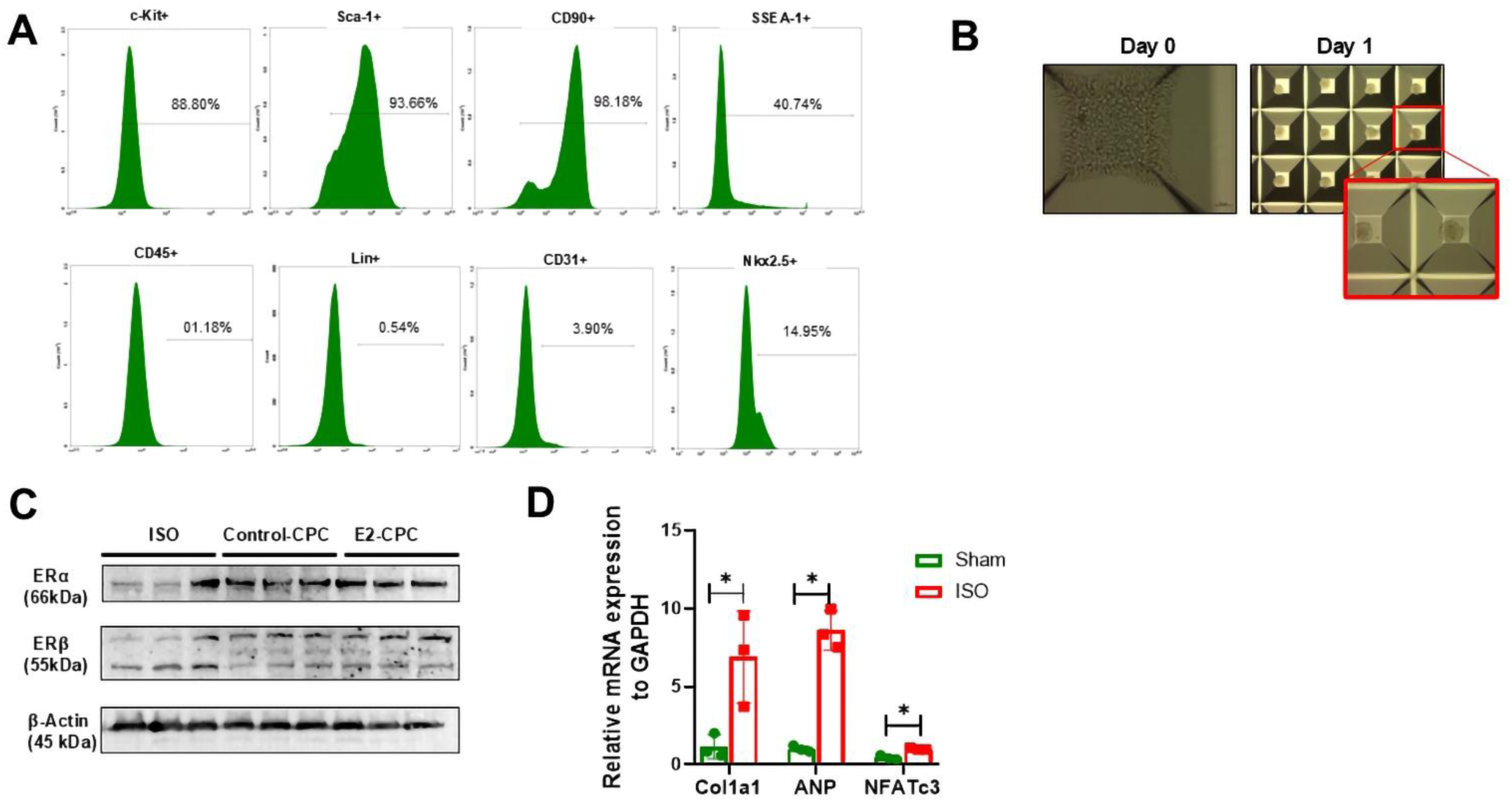
(A) Characterization of CPCs by flow cytometry analysis. **(B)** Spheroid formation from CPCs seedod on AggreWells. Images are taken by 40x, 4x and 10x from right to left,respectively. Bar represents 200 µM. **(C)** ERα and ERβ protein expression in Control-CPC and E2-CPCs was demonstrated by westen blotting.β-Actin was used as loading control. **(D)** mRNA levels of Col1a1, ANP and NFATc3 genes by qRT-PCR analysis in female mouse hearts with ISO-induced heart failure and healthy Sham control group. n=3 (biological replicate). Data are presented as mean±SD. *P < 0.05 vs Sham

**Suppl. Figure 2.**
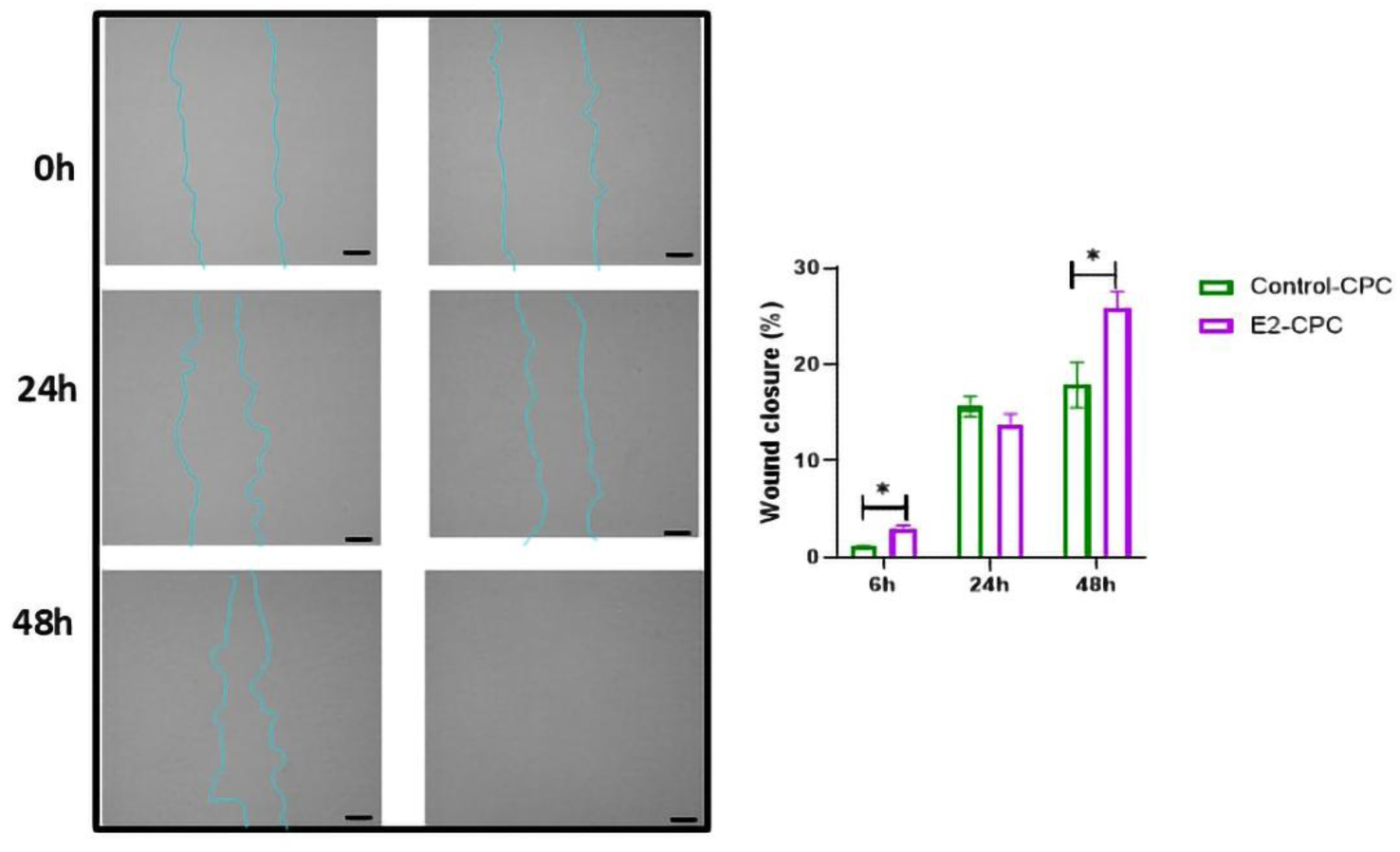
Changes in migration capacity of CPCs upon estrogen treatment was evaluated by Wound healing assay in Control-CPC and E2-CPCs groups. Images are taken at 6 h, 12 h and 24 h and by 10x. The black scale bars represent 50µM. n=3 (biological replicate). Data are presented as mean±SD.*P < 0.05 vs Control-CPC.

**Suppl. Figure 3.**
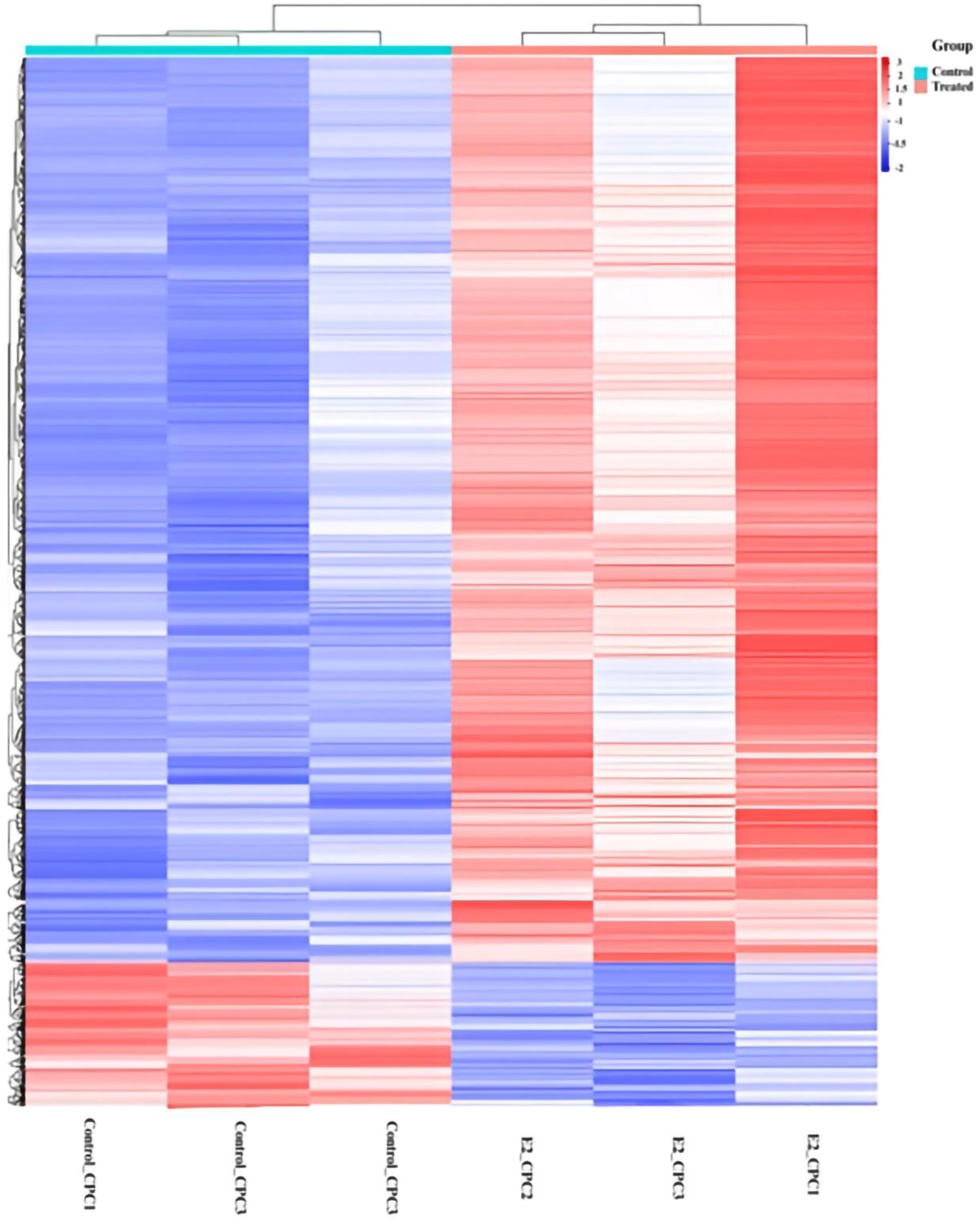
Top 1000 up and down regulated DEGs represented through heat map of the CControl-CPC and E2-CPC groups, where light blue represents the normal control group, pink represents the estogen-treated group.Red signifyes high expression and blue represents low expression.

**Table S1:**
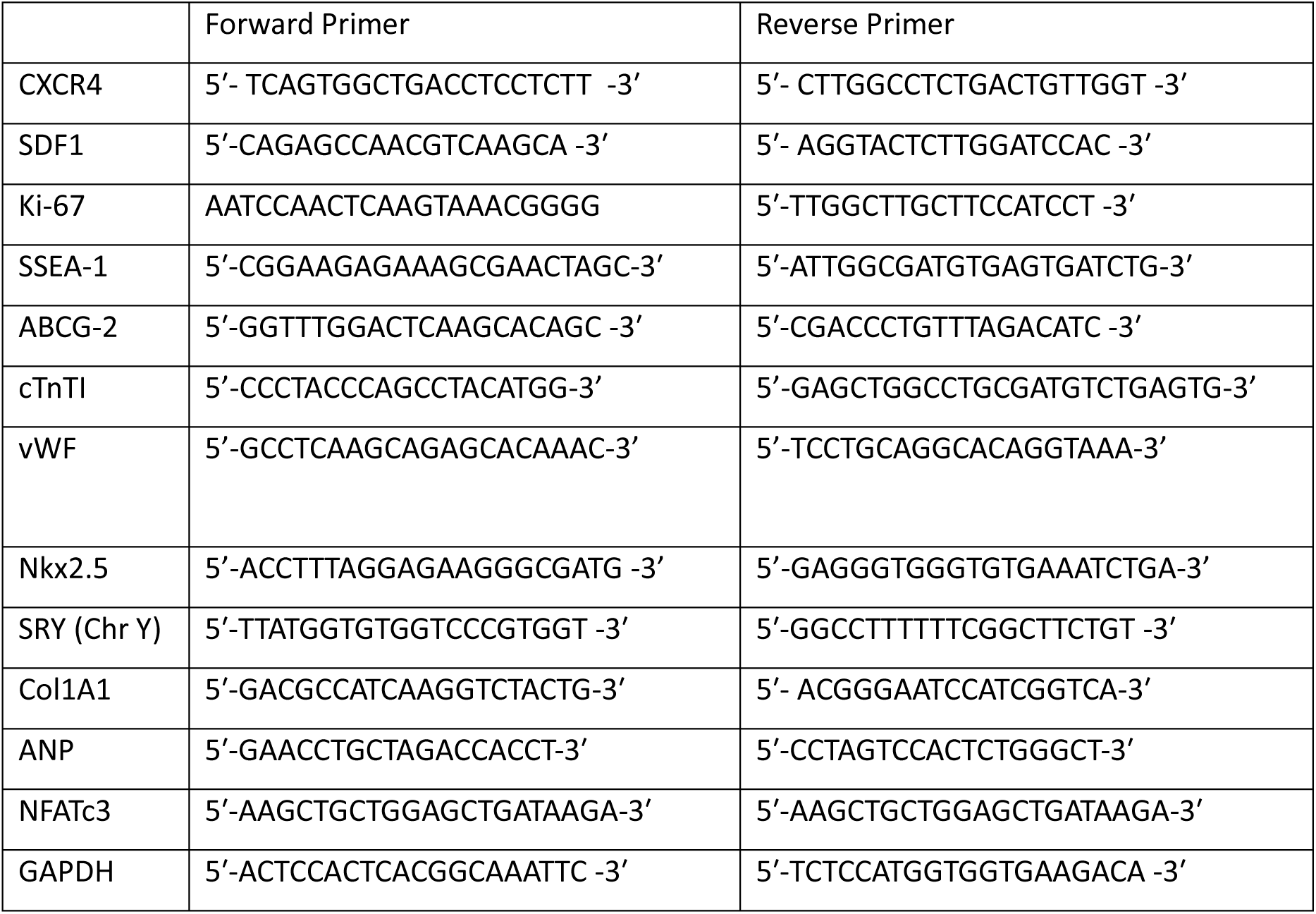
Primer sequences used for qRT-PCR.

## Notes

https://www.ncbi.nlm.nih.gov/bioproject/1242725.

### Competing Interest Statement

The authors have declared no competing interest.

